# A Structural Proteome Screen Identifies Protein Mimicry in Host-Microbe Systems

**DOI:** 10.1101/2024.04.10.588793

**Authors:** Gabriel Penunuri, Pingting Wang, Russell Corbett-Detig, Shelbi L Russell

## Abstract

Host-microbe systems are evolutionary niches that produce coevolved biological interactions and are a key component of global health. However, these systems have historically been a difficult field of biological research due to their experimental intractability. Impactful advances in global health will be obtained by leveraging *in silico* screens to identify genes involved in mediating interspecific interactions. These predictions will progress our understanding of these systems and lay the groundwork for future *in vitro* and *in vivo* experiments and bioengineering projects. A driver of host-manipulation and intracellular survival utilized by host-associated microbes is molecular mimicry, a critical mechanism that can occur at any level from DNA to protein structures. We applied protein structure prediction and alignment tools to explore host-associated bacterial structural proteomes for examples of protein structure mimicry. By leveraging the *Legionella pneumophila* proteome and its many known structural mimics, we developed and validated a screen that can be applied to virtually any host-microbe system to uncover signals of protein mimicry. These mimics represent candidate proteins that mediate host interactions in microbial proteomes. We successfully applied this screen to other microbes with demonstrated effects on global health, *Helicobacter pylori* and *Wolbachia*, identifying protein mimic candidates in each proteome. We discuss the roles these candidates may play in important *Wolbachia*-induced phenotypes and show that *Wobachia* infection can partially rescue the loss of one of these factors. This work demonstrates how a genome-wide screen for candidates of host-manipulation and intracellular survival offers an opportunity to identify functionally important genes in host-microbe systems.

## Introduction

The intracellular niche of symbiotic and parasitic bacteria is a complex environment and exhibits pressures that lead to unique coevolutionary products such as molecular mimics^1^. Intimate cellular and subcellular interactions between hosts and symbionts can create conditions in which it is advantageous for a symbiont species to produce a molecule or protein that mimics the structure of a host component. For example, molecular mimicry is a major contributor to host immune system evasion and microbe-host and virus-host coevolution ^2–6^. Distant structural homology and convergence with host proteins is likely a key evolutionary step in the generation of successful host-microbe interactions. However, successfully identifying and characterizing protein mimics often requires painstaking molecular approaches. Therefore, novel approaches for rapidly and inexpensively identifying candidates will enable fundamentally new insights and analyses.

Protein mimicry is often assessed in the context of host immune evasion for enacting control of host cellular machinery. Protein mimics have been a research focus of interactions between *Legionella* and its human host^2^, where bacterial protein mimicry is thought to contribute to manipulating numerous host systems such as transcription, ubiquitination, and protein phosphorylation. ^5,7,8^ We thus have a breadth of wet lab validation of known protein mimics that can be used to confirm and control a novel screen in a system with little structural, and biochemical data available. The *Legionella pneumophila-*human system provides a framework that bounds the extent of structural conservation within host-microbe alignments that would be of interest to identify. *Legionella*-human structural proteomes also provide a type of ‘positive control’ for any approach to identify protein mimicry.

Structural comparison allows for high sensitivity and throughput, which was limited in previous approaches to protein mimicry identification. Sequence based screens for protein mimicry have been successful in identifying candidates, but are limited in their sensitivity because they require high sequence similarity.^9^ Hebert *et al*. utilized RNA-seq data of protein coding genes and ORF predictions to construct a pipeline that identified similar protein sequences between host and symbiont genomes^6^. Other sequence based screens have also been developed for the purposes of identifying novel *L. pneumophila* effectors. These implement a variety of wet-lab and *in silico* techniques and leverage the large set of known, validated effectors^10,11^. However, while protein sequence analysis is a useful approach for uncovering eukaryotic domain-containing mimics, it disregards newly available structural data that we can now leverage for much more sensitive homology screening. Furthermore, sequence-based approaches miss convergent structural and functional mimics that may have virtually no sequence similarity. While wet lab structural and biochemical data is still the gold standard for confirming and characterizing protein mimicry, novel structure-aware approaches are needed to identify diverse mimic candidates in genetically intractable organisms.

Modern advances in protein folding prediction from genomic sequences has enabled *ab initio* assays for protein structural mimicry between symbiont proteomes and those of their hosts for the first time. Intracellular *Wolbachia pipientis* alphaproteobacterial symbionts of diverse arthropod and nematode species are ideal candidates for such a mimicry screen. *Wolbachia* genomes are enriched in eukaryotic-like elements ^12^, they induce a range of unique host-associated phenotypes, and the wMel strain from *Drosophila melanogaster* is important in applied mosquito biological control strategies.^13^ However, the functions of only five of *w*Mel *Wolbachia’s* ∼1200 genes have been directly experimentally determined because the obligate intracellular lifestyle of these bacteria has made them difficult to characterize genetically.^14–17^ Thus, a genome-wide structural protein mimicry screen offers an appealing opportunity to identify functionally important *Wolbachia* genes for interacting with and controlling host biology that could be applied in biological control strategies. The model *D. melanogaster* system naturally infected with the *w*Mel strain offers an experimental system to begin to test and validate these candidates.

Deep learning and generative AI have revolutionized our ability to predict protein structure from amino acid sequence data^18–20^, and open powerful new avenues for identifying structural similarity. Protein structure prediction models such as AlphaFold2^21^ coupled with access to large graphical compute power through cloud resources have enabled small budget access to entire structural proteomes in a matter of days. In the first half of 2022, folded proteomes were made even more accessible with the expansion of the AlphaFold structural databases, releasing predictions for nearly every sequenced organism in all domains of life. Ultrafast structural alignment methods provide tools for grappling with these large amounts of structural data. The 3Di+AA method core to Foldseek^22^ has proved to be a lightning fast means of aligning thousands of protein structures and was tested with remote protein homology searches in mind. These new approaches therefore offer an appealing opportunity to investigate molecular mimicry between symbiont and hosts genome-wide.

Here, we utilize these new genomic datasets, structure predictions, and rapid alignment tools to search for distant protein structural similarity between entire host and symbiont proteomes. We initially develop this screen utilizing the extensive knowledge of structural mimicry in the *Legionella*-human system for validation and initial testing. We then apply the validated framework to *Helicobacter pylori*, which imparts numerous host control and manipulation mechanisms on human gastric epithelial cells that lead to disease. We next apply our framework to *Wolbachia melanogaster* which infects *Drosophila melanogaster* as the host genome is extensively annotated and we have *in vitro* and *in vivo* system for potential follow up experiments on identified mimicry candidates^23,24^. These platforms allow us to establish functional assays that support the findings of the protein mimic screen, demonstrating a basis for a *Wolbachia* wMel mimic of the *Drosophila* protein, eato. We refined the structural homology hits from this process to ensure that the mimicry candidate identified is most likely a result of convergent evolution, opposed to deep phylogenetic conservation for endogenous housekeeping functions. These candidates represent unique novel connections between hosts and their microbes as well as co-evolved products resulting from a symbiotic evolutionary history. This approach advances bioinformatic investigations into convergent evolution within host microbe systems, and produces novel candidate genes for researchers seeking to perturb symbiotic systems.

## Methods

### AlphaFold Protein Structure Predictions

We downloaded predicted structural proteomes for a selection of host genomes, host-associated bacterial genomes, and free-living bacterial genomes. We selected host proteomes for *Drosophila melanogaster* and human and proteomes for their associated microbes: the wMel strain of *Wolbachia, H. pylori 26695* and the Philadelphia 1 strain of *L. pneumophila*. To control for similar structures constrained by housekeeping functions, we randomly sampled 104 free-living bacteria from across the bacterial tree of life.

### Free-Living Proteome Control Database Generation

The free-living control method described above was developed with the goal of ensuring alignments between host and microbes were unique to those systems and not representative of a highly conserved structure found within most bacteria. Highly conserved symbiont proteins required for housekeeping functions could exhibit structural similarity with host proteins due to shared endogenous functions. These proteins would be highly similar to free-living orthologs, given their close phylogenetic proximity and necessity for core cellular processes, allowing us to identify and exclude them from the mimic candidate gene set. Leveraging this structural conservation logic, we developed a method that would confidently ensure that each candidate mimic protein was not core to free-living bacterial cellular function, and instead had evolved a divergent function through converging on a common structure with its host.

In order to develop a large enough control proteome dataset that would sufficiently describe free-living proteomes and their structural content, we automated the selection process. Specifically, we designed a system of randomly selecting bacterial species and automating the literature search to determine if the selected species was free-living or potentially host-associated. The motivation for this method for developing such a dataset is in part due to the lack of annotation on host-association of bacteria in current databases. We utilized taxonKit^25^ to generate a json representation of the entire NCBI bacterial taxonomy database. We then used a custom python script to randomly search this dataset structure by bacterial order and species. To do this we randomly selected two species from each randomly selected order. The script uses the NCBI Entrezpy python library to validate that a selected species has a genome in the assembly database that is marked as complete. Randomly selected bacteria species were then processed with another custom python script leveraging the unofficial Bard-API (https://github.com/dsdanielpark/Bard-API) accessed on October 20th, 2023 and the BacDive database API^26^. Literature related to the bacteria species queried was processed by a constrained bard session in order to make conservative predictions on whether or not the bacteria was free-living or potentially host-associated. We manually reviewed these selected taxonomy IDs and used them to download associated proteome “shards” from the AlphaFold databases. Free-living proteomes that contained at least 900 structures were selected for use in the control dataset. The full list of organisms selected for this database along with the literature used in the decision is available in the supplementary data (Supplemental Table S1).

### Protein Structure Alignments

We performed structural alignments between proteomes with the tool Foldseek^22^. Foldseek discretizes structures into sequences that describe the spatial relationships between neighboring residues. We selected the 3Di + AA local alignment method for its superior alignment speed while retaining similar sensitivity to the global TM-Align method ^22,27^. We treated microbe proteins as queries and host proteins as targets, generating a TM-score ranging from 0 to 1 for primarily evaluating the structural similarity of an alignment pair. We selected an e-value cutoff of 0.01 for these alignments, an established metric used in large scale alignment and clustering methods developed by the Steinegger lab^28^.

### Structure Alignment Controlling and Filtering

We designed and tested our methods with the intent to filter out proteins that are conserved structures in free-living bacteria, and thus likely not examples of host manipulation or symbiosis. We first filtered alignments for mimic candidates by their TM-score, target coverage, query coverage, fraction of identical residues and fraction of free-living proteome alignments. A high fraction of identical residues in alignments is evidence of recent shared ancestry, possibly due to horizontal transfer. This value is expected to be large for recently transferred genes, or highly conserved housekeeping genes^29^. A high fraction of free-living proteomes containing a significant alignment with a query protein indicated the overall conservation of the structure in free-living proteomes. This may suggest that the microbe protein is less likely to be involved in host interaction or manipulation. In order to further filter the resulting list of candidates to arrive at a final list of high quality protein mimicry candidates, we also considered qualitative metrics such as gene ontology and conservation of critical residues and domains within the structural alignment.

We obtained protein mimic positive control sets for *Legionella* from Mondino *et al* 2020^2^. This set of 23 genes represented validated *Legionella* effectors showing structural mimicry. *Legionella* proteins were classified as eukaryotic domain containing or eukaryotic like proteins which we equate with partial and full mimicry, respectively. Alignments between this positive control set and the human proteome were used to determine reasonable filters for alignment statistics. For controlling alignments we removed non-host-associated structural conservation based on alignment to the free-living bacterial proteomes dataset. We aligned the host-associated microbial proteome to each of the proteomes in this dataset, allowing us to report a fraction of proteomes that the microbe protein had a reciprocal best hit alignment within.

### GO Enrichment Analysis

We used host protein UniProt IDs and the PANTHER^30^ classification system for the GO enrichment analysis^31,32^. P-values were obtained with Fisher’s exact test. We used FDR for correction.

### Phylogenetic Tree Construction

We performed a BLAST+ 2.15.0^33^ search with the *Wolbachia* wMel *HtpG* gene sequence (WP_022626167.1) and manually selected samples from the results retaining the amino acid sequences for each hit. We next used ClustalO v1.2.4 to align the amino acid sequences selected from the blast results and manually trimmed the core alignment to 923 amino acid positions. We used IQTree v2.2.5 with standard amino acid tree inference parameters and 100 bootstrap replicates. This consensus tree was plotted in FigTree with further annotation in Inkscape.

### Ovary fixation and PI staining

We derived our methods from previous work done on the *Drosophila* oocyte^23^. Newly enclosed flies were transferred to fresh white food with yeast and aged 4–5 days. 10–15 flies from each cross were dissected, fixed, and stained with propidium iodide (PI). Ovaries were dissected, the ovarioles were separated with pins, and fixed in 200 μl devitellinizing solution (2% paraformaldehyde and 0.5% v/v NP40 in 1x PBS) mixed with 600 μl heptane for 20 min at room temperature, with agitation. Oocytes were then washed 5x in PBS-T (0.1% Triton X-100 in 1x PBS), and treated with RNAse A (10 mg/ml) overnight at room temperature. Oocytes were washed six times in PBS-T and were incubated overnight in PI mounting media (20 μg/ml in 70% glycerol and 1x PBS), and mounted on glass slides. The *uninfected OreR* stocks were used as wild-type controls. Uninfected *Eato* mutant and wMel-infected *Eato* mutant samples were processed simultaneously to minimize batch effects. Wild-type control samples were processed after these experimental samples. Slides were imaged immediately or stored at -20°C for no more than three weeks. Oocyte localization variation is the same between runs for wild-type and mutant genotypes.

### Confocal imaging

Stage 14 egg chambers were imaged on a Leica SP5 confocal microscope with a 63x objective. Optical sections were taken at the Nyquist value for the objective, every 0.34 or 0.38 μm, at a magnification of 1x. Propidium iodide was excited with the 514 and 543 nm lasers, and emission from 550 to 680 nm was collected. Images were processed in ImageJ^34^.

### Image and plotting analysis

To quantify engulfment, we counted the number of persisting NCs in each stage 14 egg chamber through z-stack and 3D reconstruction on ImageJ. The criteria for a stage 14 egg chamber was fully developed dorsal appendages^35^. For better visualization on ImageJ, the brightness/contrast tool was used to increase the threshold on the image, and all background noise was rendered black. The resulting NC counts for all genotypes were plotted in R through ggplot2^36^. The resulting NC counts between uninfected and wMel infected *Eato* mutants were compared with the Wilcoxon Rank Sum Test.

## Results and Discussion

### Establishing Expectations for Protein Mimic Alignments using Known Legionella Examples

We leveraged well-characterized examples of *Legionella* mimicry of human proteins and a large database of free-living proteomes to establish our baseline expectations for genome-wide candidate structural mimicry detection. First, we examined the expected TM-Scores, target coverage, query coverage, fraction of identical residues and fraction of free-living proteome alignments of protein structure alignments using a subset of 14 biochemically characterized protein mimics from *Legionella*-human examples (Supplemental Table S2). We also considered their classification according to Mondino *et al* 2020, which categorizes effectors into eukaryotic domain containing and eukaryotic resembling proteins. We referred to these classifications here as partial and full mimics. Then, we generated a structural proteome alignment between the human proteome and *Legionella pneumophila* subsp. *pneumophila* str. Philadelphia 1 to use as a validation set for investigating further host-microbe pairs *ab initio*.

Analysis of the validation mimic set for *Legionella* confirms that our screen can robustly detect both full and partial structural protein mimics. We found that TM-Scores of the known eukaryotic domain-containing *Legionella* proteins ranged from 0.4-0.8, with an average of 0.64, target (host) coverage ranged from 0.1-0.95, with an average of 0.43 and query (microbe) coverage ranged from 0.23-0.99, with an average of 0.67. This large target coverage range tightened significantly for the full structural mimics to 0.6-0.97 with an average of 0.82 while the TM-Score range for full mimics encompassed values from 0.4-0.95 with an average of 0.65. We found that the fraction of free-living proteomes aligned differed greatly between *Legionella’s* 5 known full structural mimics and the 9 partial ones with averages of 0.6 and 0.2, respectively.

Overall this validation informed our expectations on the search for structural mimicry to require generous thresholding around the different expected ranges for the two general classifications of mimicry extent. This was necessary in order to encompass all possible examples of mimicry in our alignment results. Using an E-value cutoff of 0.01, we decided to select results for further ontological evaluation with structural statistical thresholds of 0.4 for TM-score and 0.25 for target coverage. We also determined there could be cases where partial mimicry of a eukaryotic domain could occur within a large host protein. In order to catch these cases we did not filter alignments with lower than 0.25 target coverage if the query coverage was at least 0.5. In the case of our *Legionella* and Human proteome alignment ∼35,000 of the 45,752,796 alignments performed met these initial filters. After filtering based on TM-score, target and query coverage we still retained 5,227 unique microbe proteins participating in thousands of alignments (Supplemental Table S3). Our controlling method based on the fraction of aligned free-living proteomes further reduced the total number of alignments considered. 2600 alignments exhibited free-living frequencies equal to or less than the average for partial mimics in *Legionella* of 0.2.

### Comparing Host-Microbe Structural Proteomes Identifies Candidate Mimics

We applied our framework to three host-microbe systems: *Legionella-*human, *H*.*pylori-*human and *Wolbachia* wMel-*Drosophila melanogaster*, identifying strong candidate mimic-like effectors in each. Comparing the overall trends between microbe-host and microbe-free-living bacteria alignments revealed key differences likely attributable to differences in gene content between *w*Mel, *H. pylori*, and *Legionella* (Figure 1 and Figure S1). Consistent with its larger genome (3.5 Mb vs 1.3 Mb and 1.6 Mb), *Legionella* encodes more proteins with significant structural alignments to its host genome than *Wolbachia* or *H. pylori* encode. We noticed that *Wolbachia* and *H. pylori* appeared to have a higher fraction of free-living conserved proteins than *Legionella*, but found that the total number of proteins highly conserved in free-living proteomes between the three organisms was comparable. At a fraction of free-living proteomes aligned cutoff of 0.8, the number of query structures present in free-living proteomes from *Wolbachia* and *H. pylori* was 297 and 364 respectively. In *Legionella* this value was 384, despite *Legionella* having twice the number of protein structures. The difference between the conservation statistics between these endosymbiotic bacteria is likely due to the nature of host association in these systems. In the case of *Wolbachia*, strains verge on mutualism with many of their various hosts^24,37,38^. *H. pylori* infects a narrow range of hosts and is highly adapted for human gastric mucosa^39,40^. In both these cases the highly specified nature of association with a host has led to a more streamlined genome. In contrast, *Legionella* is a damaging pathogen driven by a large genome overrun by elements from many previous eukaryotic hosts that it has evolved within^41^. However all of these systems have protein structures that are rarely found within free-living bacteria from which we have selected strong candidates for protein mimicry of host cellular machinery.

**Figure 1.**
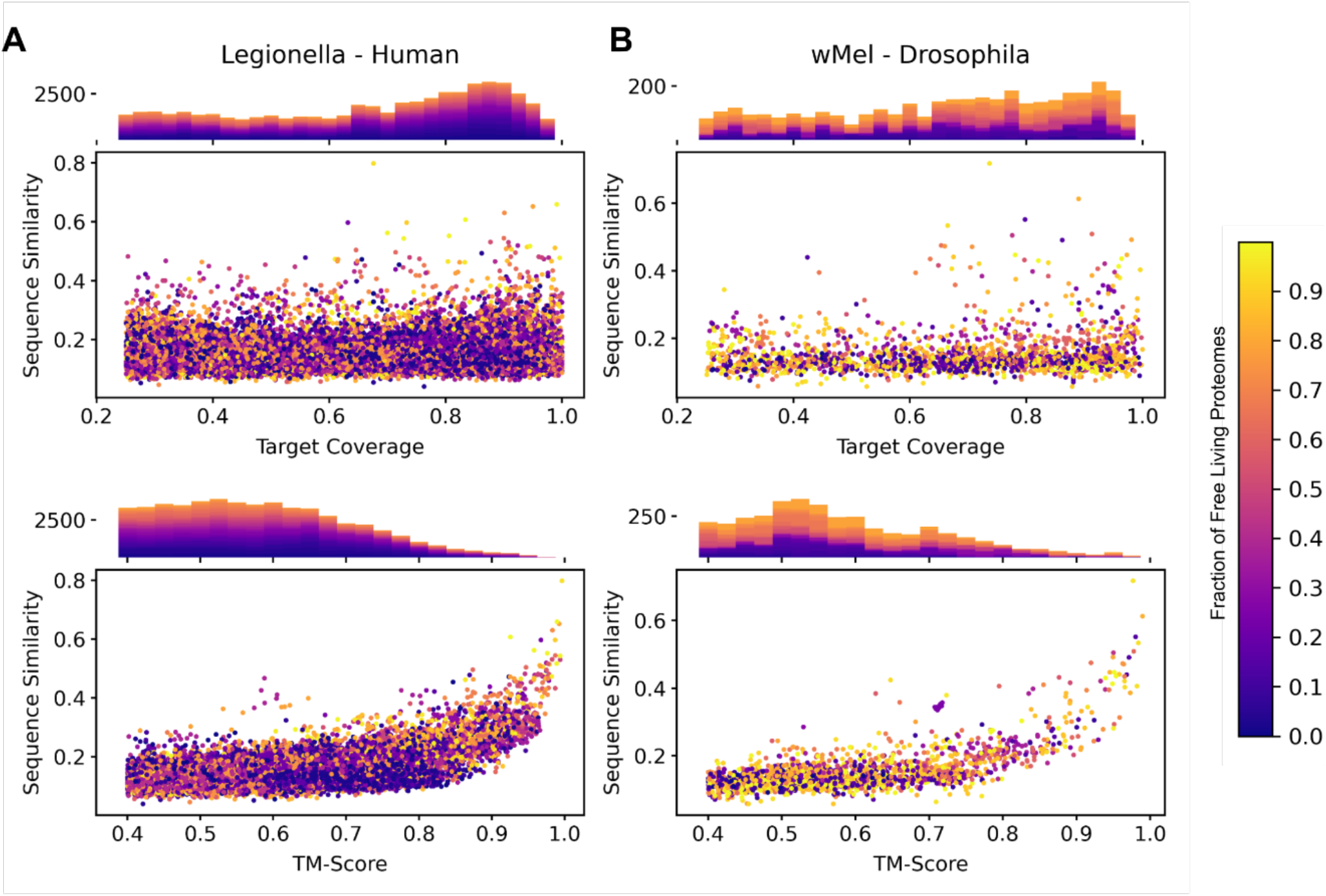
Alignment Space Under Expansive Control Method. **A**. *Legionella*-Human alignment space sequence similarity as it relates to target coverage and TM-Score. The averages for the fraction of free-living proteomes aligned for both systems were 0.416 for *Legionella-*Human and 0.605 for wMel-*Drosophila*. **B**. wMel-*Drosophila* alignment space sequence similarity as it relates to target coverage and TM-Score. Alignments are colored by the fraction of control proteins the query protein was aligned with.

### Structural Mimicry in *Legionella pneumophila*

Structural mimicry of both partial and full host proteins is a common method of host cell manipulation used by *Legionella pneumophila* strains, and is why we selected this pathogen for validation of this screen. *L. pneumophila* is a Gram-negative bacterium and the causative agent of Legionnaires’ disease, a severe form of pneumonia. *L. pneumophila* strains, like other bacterial pathogens, must avoid phagosome fusion with lysosomes through evolution of numerous host cell manipulation processes^42^. A vast arsenal of host effectors and mimics has been painstakingly uncovered through years of biochemical research^2^. However, *L. pneumophila’*s bloated pathogenic genome is influenced by countless eukaryotic host associations through which it acquired mobile genetic elements and genes by HGT, and is likely to still have mimic-like effectors yet to be uncovered^41^. We also performed a GO analysis of the host target proteins aligned to the 2,600 microbe protein subset obtained through filtering based on the free-living proteome alignment fractions discussed in the validation section above (Supplemental Table S4). Many of the top enriched biological processes were associated with transmembrane transport. It is important to note that bacterial queries were not limited to alignments with a single host target structure and single query structures contributed multiple targets to the protein IDs used in the GO analysis. Hence we did not rely on GO analysis results for finding strong examples in the data, and rather noted the results as potentially informative of mimic trends. In our use of *L. pneumophila* as a validation organism for our framework, we were able to identify strong candidates for mimic-like effectors previously uncharacterized. In addition to these effector protein candidates, we were able to suggest new interactions and mechanisms of known effectors, such as Mip, through reconsidering them as mimic-like as they appeared in our results.

#### Q5F2U4 (Mip)

Mip is a well known *L. pneumophila* effector proven to be essential for virulence in various host systems^43^. While Mip has been previously characterized, it has not been conclusively determined what components of human cells Mip interacts with. A recent paper showed that Mip has a general interaction with and regulates the lncRNA GAS5/miR-21/SOCS6 axis influencing phagocytosis and chemotaxis in the RAW264.7 macrophage-like cell line^44^. Similarly it has also been shown that *L. pneumophila* infections dampen the cytokine response of infected macrophages^45^. The *L. pneumophila* structure Q5F2U4 aligns with a human peptidyl-prolyl cis-trans isomerase, FKBP1A which has been shown to have a binary interaction with suppressor of cytokine signaling 6 (SOCS6)^46^. These protein structures aligned with a TM-score of 0.89, target coverage of 0.68, query coverage of 0.4, and fraction of identical residues of 0.36 (Figure 2). Thus, this potential example of mimicry supports previous predictions about Mip interactions with host cellular machinery, and suggests a mechanism for interaction in the mimicry of FKBP1A or other structurally similar host peptidyl-prolyl cis-trans isomerases involved in cytokine suppression.

**Figure 2.**
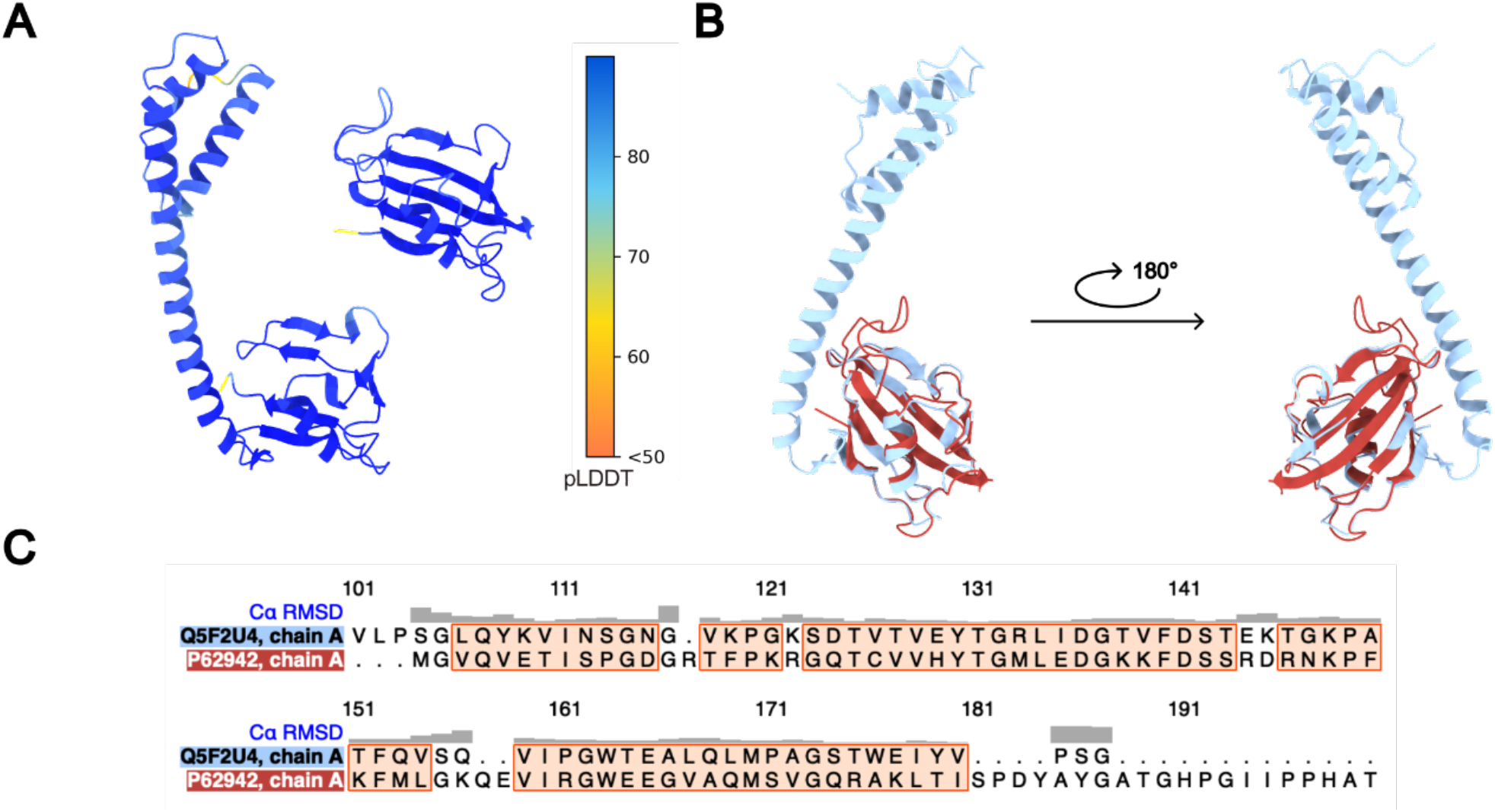
Mimic Candidate Q5F2U4. **A**. Structures of *L. pneumophilia* Q5F2U4 and human FKBP1A colored by pLDDT. **B**. Structural alignment of *L. pneumophila* (blue) and human (red) proteins. **C**. Portion of sequence alignment between microbe and host proteins, showing low sequence similarity. Similarity between the two structures is shown by the RMSD at the top of the plot.

#### Q5ZW23

This *L. pneumophila* protein is predicted to be a membrane associated proton/peptide symporter and it aligns to the full human Solute Carrier Family 15 member 3 (SLC15A3) protein. It shares the strongest alignment with Q8IY34, SLC15A3 (TM-score: 0.73, target coverage: 0.91, query coverage: 0.98 fraction identical residues: 0.15, Supplemental Figure S2). The structure is not found in any of the free living proteomes screened. The human protein is associated with the lysosomal membrane^47^ and is thought to be involved in the detection of microbial pathogens by toll-like receptors and NOD-like receptors through studies in mice^48^. This example of potential mimicry could thus coincide with *L. pneumophila’*s ability to avoid fusion with the lysosome and represent another tool in its extensive kit of subverting host immune responses.

### Structural Mimicry in Helicobacter Pylori

Other host-associated bacteria that are limited in their experimental tractability can be investigated with our approach, including human pathogens such as *H. pylori*. The extracellular bacterium infects the stomach epithelium and the upper part of the small intestine in humans^49,50^, and has also been found in primates, rodents and other various mammals^51^. *H. pylori* can cause various gastrointestinal malignancies, including inflammation of the stomach lining and ulcers. *H. pylori* infection is quite common, estimated to infect about half of the world’s population^49^. Its culture requires a microaerophilic environment and complex blood-containing media, making it challenging to study *in vitro*^52^. In order to establish such persistent and widespread infections, *H. pylori* utilizes highly specialized mechanisms to avoid host defense and adaptive immune mechanisms in gastric epithelial cells. The bacteria interact closely with host cells, leading to manipulation of their cellular process and functions^53^. These interactions cause a wide range of effects including DNA damage, apoptosis, proliferation, stimulated cytokine production and even cell type transformations^53,54^. Although dozens of effectors secreted by *H. pylori’*s type IV secretion system (T4SS) have been identified, many more effectors remain to be discovered^50,55,56^.

We produced 24,139,512 pairwise alignments between the *H. pylori* strain 26695 and human proteomes. 3516 unique microbe protein structures had alignments to host proteins that met our significant alignment criteria (Supplemental Table S5). Of these, 888 were depleted among free-living proteomes (aligned to a fraction less than 0.2). We performed a GO analysis of the host target proteins aligned to this 888 microbe protein subset (Supplemental Table S6). Many of the top hits were associated with cellular transport such as asparagine, leukotriene, (R) - carnitine and leucine transport into and between cells. Similar to our screen of the *Legionella* proteome, the application to *H. pylori* identified both novel candidates for structural mimicry as well as suggested mechanisms for already existing effector candidates.

#### O25525 (guaC)

The *H. pylori* O25525 protein is a GMP reductase responsible for catalyzing NADPH-dependent deamination of GMP to IMP in purine salvaging and processing pathways, which aligned with the full structure of a human GMP reductase. *H. pylori* is reliant on deamination pathways for purine nucleotide biosynthesis, as it lacks the enzymatic machinery necessary for *de novo* production of IMP and must salvage purines from the environment^57^. A study into bacterial mutants lacking each of the critical components of the pathway have shown varying capabilities to utilize different purine bases and nucleosides, with guaC mutants having limited conditions for growth^58^. *H. pylori* has a strong, full structural alignment with the human GMP reductase 2 (GMPR2) protein, with a TM-score of 0.876 a target coverage of 0.98 and a fraction of identical residues of 0.32 (Supplemental Figure S3). O25525 was nearly non-existent in the free-living control set with an alignment fraction of 0.02. It has been shown that GMPR2 overexpression can promote differentiation of leukemia cells^59^. This could suggest that *H. pylori* expression of a highly similar structure could be promoting gastric cancer development.

#### O25981

A periplasmic protein of unknown function is encoded by the *H. pylori* gene *HP1440*, which exhibits full structural alignment with a human serine protease. *HP1440* has been recently shown to belong to the CrdRS regulon^60^. The CrdRS regulon is a set of genes controlled by the CrdRS two-component system in response to changes in its environment, specifically in the context of host-induced stress. *H. pylori* O25981 aligned with the human chymase CMA1 structure P23946, with a TM-score of 0.49, target coverage of 0.99, query coverage of 0.96, and fraction of identical residues of 0.11 (Figure 3). This serine protease is expressed in mast cells where it activates Matrix metalloproteinase (MMP)-9 and induces inflammation^61^. In *H. pylori* infections, MMP-9 gene expression is significantly increased after *H. pylori* infection, potentially by activation with its own chymase-mimic^62^.

**Figure 3.**
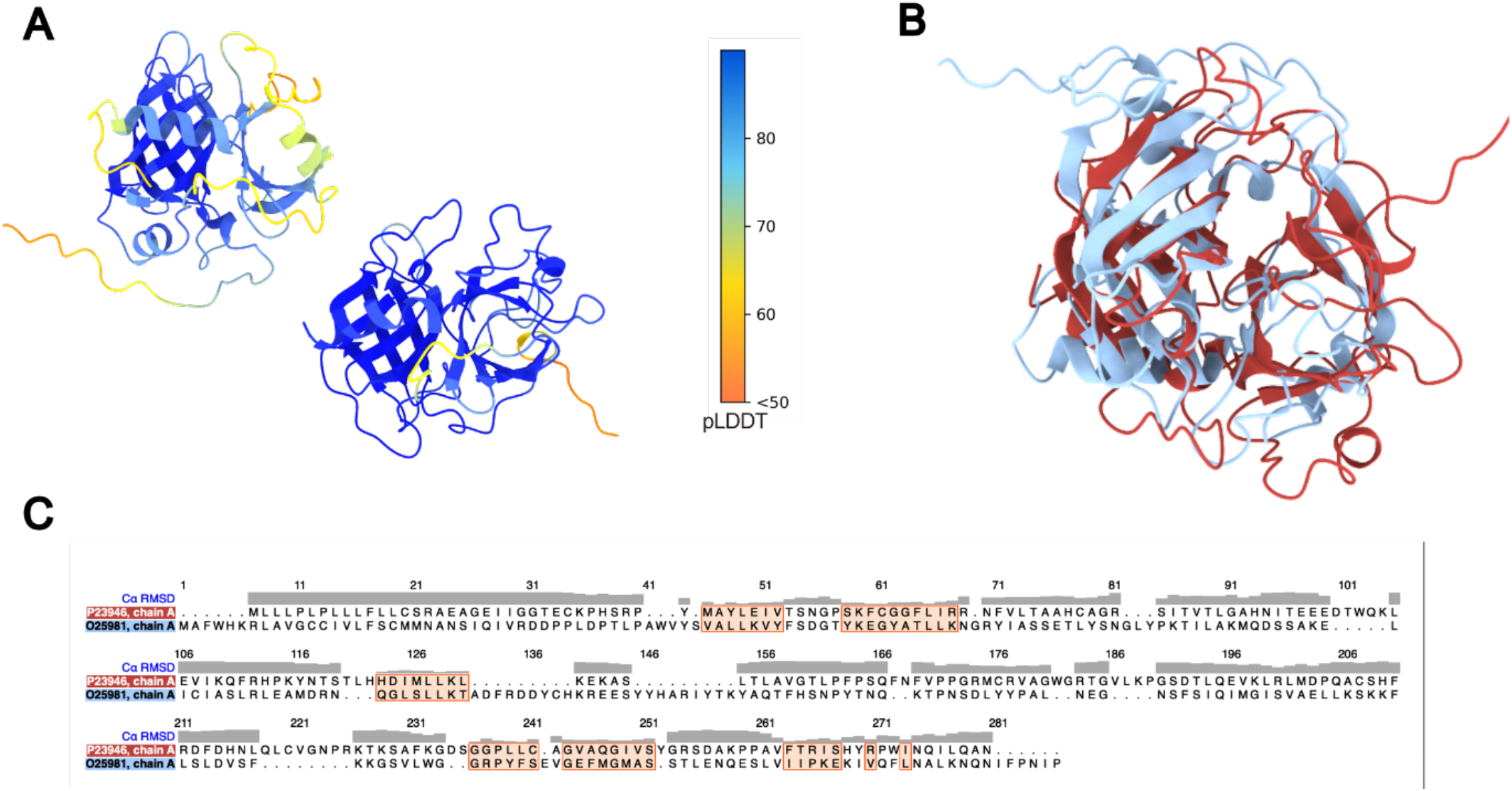
Mimic Candidate O25981. **A**. Structures of *H. pylori* O25981 and human P23946 colored by pLDDT. Structures exclude extension of terminal disorder tails. **B**. Structural alignment of *H. pylori* (blue) and human (red) proteins. **C**. Sequence alignment between microbes and host proteins showing low sequence similarity. Structural alignment similarity is shown by the RMSD at the top of the plot.

### Structural Mimicry in Wolbachia wMel

We performed ∼13,700,000 pairwise structural alignments between the *wMel* and *Drosophila* proteomes. From this full proteome alignment, 4,644 alignments from 460 unique *wMel* protein structures fell within the predetermined range of interest in the alignment space for candidate structural mimics. Using the 0.2 threshold for filtering left 33 *wMel* proteins with significant alignments (Table 1).

**Table 1.** Top Hits in wMel-*Drosophila* Alignments. Alignments remaining after applying both filtering (TM-Score, Target Coverage, E Value) and controlling (Fraction of Free-living Proteomes Aligned) metrics to the initial set of 13,700,000 structural alignments. This table shows only the top hit for each wMel protein. The full list can be found in Table S2.

| Microbe UniProt ID | Host UniProt ID | TM-Score | Target Coverage | Query Coverage | Fraction of Identical Residues | Fraction of Freelifing Proteomes Aligned |
| --- | --- | --- | --- | --- | --- | --- |
| Q73HM8 | Q9VIE7 | 0.5032 | 0.221 | 0.798 | 0.138 | 0 |
| Q73HH1 | A1Z7R8 | 0.5999 | 0.449 | 0.782 | 0.141 | 0 |
| Q73HH7 | Q27297 | 0.6376 | 0.658 | 0.95 | 0.144 | 0 |
| Q73G80 | P18932 | 0.7179 | 0.993 | 0.987 | 0.143 | 0.009 |
| Q73HU7 | Q9VUN9 | 0.6246 | 0.452 | 0.532 | 0.134 | 0.019 |
| Q73IF6 | E1JHQ6 | 0.5214 | 0.784 | 0.989 | 0.142 | 0.028 |
| Q73I63 | P00399 | 0.7291 | 0.906 | 1 | 0.153 | 0.028 |
| Q73HH8 | O77135 | 0.5108 | 0.812 | 0.417 | 0.118 | 0.047 |
| Q73HF3 | Q7JZB4 | 0.7717 | 0.642 | 0.984 | 0.127 | 0.047 |
| Q73GY6 | Q7JXC6 | 0.4964 | 0.591 | 0.971 | 0.199 | 0.047 |
| Q73H00 | Q9VV59 | 0.7718 | 0.928 | 0.995 | 0.292 | 0.047 |
| Q73HR0 | A0A0B4K701 | 0.5631 | 0.94 | 0.732 | 0.107 | 0.056 |
| Q73GW0 | Q9VKW3 | 0.6809 | 0.886 | 0.985 | 0.184 | 0.056 |
| Q73G57 | Q9VQD7 | 0.7246 | 0.937 | 0.98 | 0.248 | 0.056 |
| Q73FP3 | Q9VW26 | 0.924 | 0.891 | 0.982 | 0.305 | 0.056 |
| Q73HM6 | Q9VMU0 | 0.6812 | 0.381 | 0.833 | 0.26 | 0.065 |
| Q73IJ0 | Q9VWI0 | 0.8592 | 0.508 | 0.968 | 0.301 | 0.075 |
| Q73I50 | Q8IR48 | 0.5723 | 0.767 | 0.878 | 0.135 | 0.084 |
| Q73G22 | Q9VKW3 | 0.6287 | 0.563 | 0.808 | 0.123 | 0.093 |
| Q73I18 | Q8SWV6 | 0.7282 | 0.474 | 0.759 | 0.223 | 0.093 |
| Q73G89 | Q9VTJ8 | 0.9682 | 0.424 | 0.943 | 0.44 | 0.103 |
| Q73I37 | Q8SWV6 | 0.7367 | 0.498 | 0.768 | 0.205 | 0.112 |
| Q73GQ6 | Q9VFF3 | 0.8519 | 0.178 | 0.802 | 0.224 | 0.112 |
| Q73IX6 | A1Z9E3 | 0.9803 | 0.798 | 1 | 0.552 | 0.112 |
| Q73HH0 | A1Z7R8 | 0.4889 | 0.676 | 0.837 | 0.117 | 0.121 |
| Q73H67 | Q9VKD3 | 0.9027 | 0.81 | 0.971 | 0.307 | 0.121 |
| Q73GB2 | Q9VNP7 | 0.6204 | 0.666 | 0.854 | 0.136 | 0.14 |
| Q73GW3 | P18932 | 0.8002 | 0.764 | 0.919 | 0.215 | 0.159 |
| Q73IF8 | Q9VBI2 | 0.5848 | 0.775 | 0.961 | 0.114 | 0.168 |
| Q73GH2 | Q9VVL7 | 0.6698 | 0.683 | 0.964 | 0.182 | 0.168 |
| Q73GW2 | Q9VRD6 | 0.5262 | 0.938 | 0.604 | 0.142 | 0.178 |
| Q73HE7 | Q9VCK6 | 0.9295 | 0.963 | 0.991 | 0.247 | 0.187 |
| Q73GP4 | P29843 | 0.7154 | 0.941 | 0.984 | 0.353 | 0.196 |
| Q05HL6 | Q9VCE6 | 0.5639 | 0.352 | 0.582 | 0.15 | 0.206 |

The protein mimicry screen and subsequent ontological analysis identified examples of both full and partial structural mimicry that corroborated hypothetical mechanisms underlying *Wolbachia* wMel-*Drosophila* interactions. We identified 1,877 *Drosophila* protein targets with alignment metrics that passed our initial filters. Of these, 279 aligned to fewer than 20% of free-living proteome query structures (Supplemental Table S7). GO enrichment analysis identified terms for numerous biological processes significantly enriched in this *Drosophila* target protein set, including symbiotic interactions such as modulation by symbiont of entry into host, regulation of biological processes involved in symbiotic interaction, movement within host, and golgi vesicle fusion to target membrane. The full table of results (Supplemental Table S8) as well as files mapping query IDs to GO terms can be found in the supplemental data. The candidate proteins we identified through alignment statistics are predicted to encode functions that are consistent with known host-manipulation phenotypes, such as cytoplasmic incompatibility (CI), and also include multiple candidates associated with intracellular processes known to be modulated through *Wolbachia* infection.

#### Q73HU7

The wMel protein Q73HU7 contains a partial structural mimic to a *Drosophila* deubiquitinase. Q73HU7 contains an ovarian tumor (OTU) domain from the OTU sub-family of deubiquitinases. OTU domains are found in Eukaryotes, viruses and some pathogenic bacteria^63^. Their biochemical function is predicted to involve serine or cysteine protease activity, although loss-of-function OTU-class pseudo deubiquitinases have been reported^64^. This protein was identified through a partial structural alignment with a *Drosophila* cysteine-type deubiquitinase Q9VUN9 at a conserved OTU domain where the target and query coverages were 0.452 and 0.532 respectively. The TM-Score of the alignment was 0.62, but from the superimposed structures it is clear that the OTU domain is conserved between the two proteins (Figure 4C). Furthermore the residues thought to be critical for OTU domain function and substrate recognition are also consistent at the sequence level between the wMel and *Drosophila* proteins (Figure 4B). It is well known that host manipulation tactics employed by pathogenic bacteria such as *Legionella* often use deubiquitinase effectors^8,65^. And although their full pathways are not explicitly understood yet, CI factors used by *Wolbachia* are known to have deubiquitinase activity ^66,67^.

**Figure 4.**
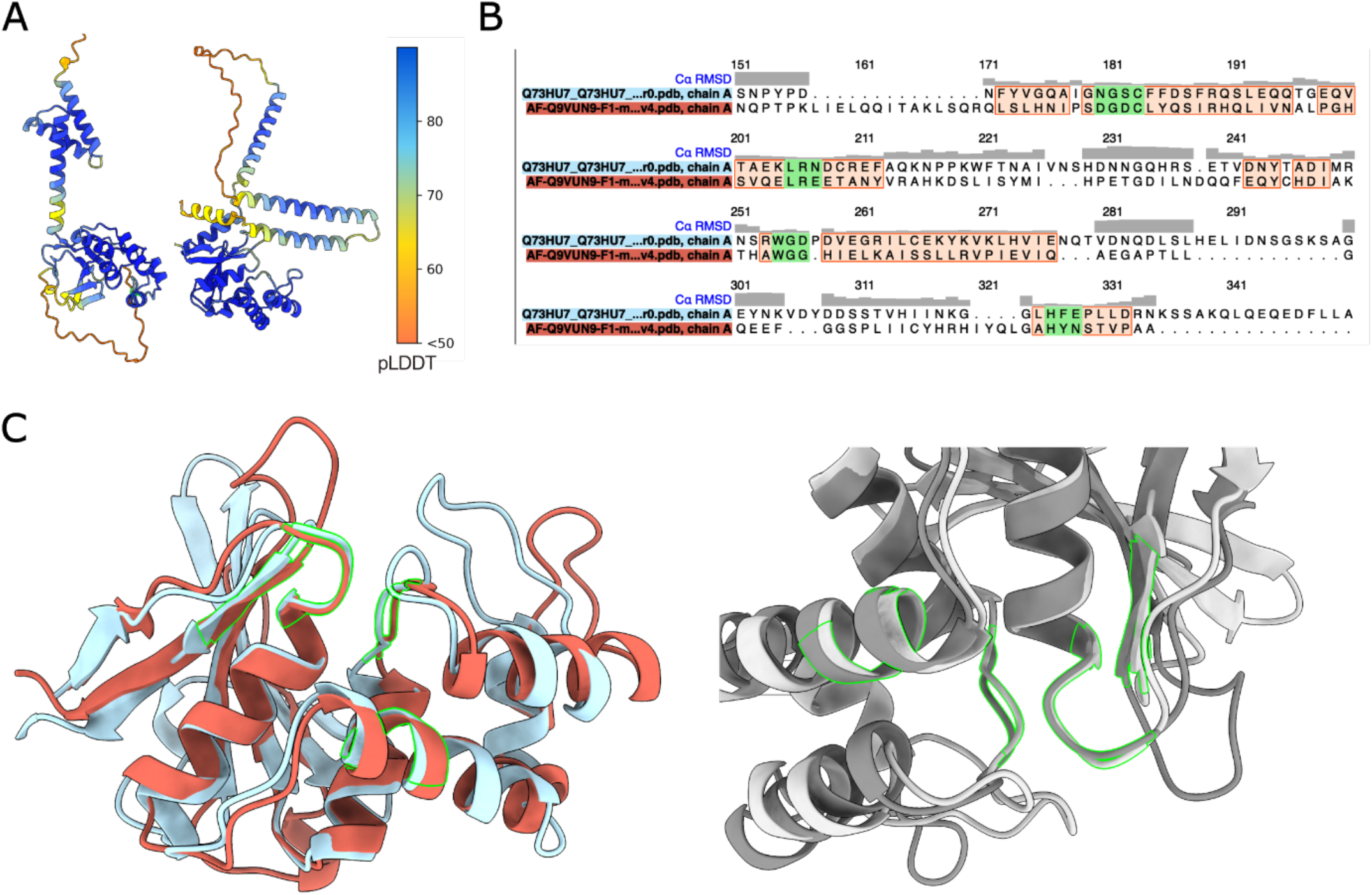
Mimic Candidate Q73HU7 Overview. **A**. Full structures of query *w*Mel *Wolbachia* Q73HU7 (left) and *D. melanogaster* target Q9VUN9 (right) colored by pLDDT showing strong model confidence in the OTU domain region. **B**. Sequence alignment at structural alignment core showing low fraction of identical residues, but high similarity of critical residues at each of the four motifs characteristic of an OTU domain. Structural alignment similarity is shown by the root mean square deviation (RMSD) at the top of the plot. **C**. Structural alignment between host and microbe proteins at the region of interest (the OTU domain). Conserved residues critical to enzyme function and substrate recognition are highlighted in green.

#### Q73IF8

The uncharacterized *Wolbachia* protein Q73IF8 aligned with the full structure of the *Drosophila* Phosphomevalonate Kinase (Pmvk) gene product, Q9VIT2 (Supplemental Figure S4). The alignment had a TM-Score of 0.53 covering 0.96 of the target structure and 0.89 of the query. The fraction of free-living proteomes aligned for Q73IF8 was 0.17. Expression of Pmvk has been shown to increase peroxisome volume in *Drosophila*^68^. Modulation of peroxisome volume and abundance may underlie the lipid metabolism modifications induced by *Wolbachia* infection, which have been attributed to influencing RNA virus replication in *Aedes albopictus* mosquito cells^69^.

#### Q73GY6

The wMel Q73GY6 RDD domain-containing protein structure partially aligned with the *Drosophila* Peroxisome assembly protein 12 (PEX12), a component of a retrotranslocation channel required for peroxisome organization. PEX12 mediates export of the Peroxisome assembly protein 5 (PEX5) receptor from peroxisomes to the cytosol^70^. The proper function of PEX12 is critical for spermatid development and inhibition of the peroxisome complex could lead to developmental defects^71^. The portion of PEX12 that the wMel protein of undetermined function aligns with is the membrane associated section, and is missing the second zinc finger domain encoded by PEX12 (Supplemental Figure S5). This deletion suggests that this protein could function as a negative regulator of proper peroxisome assembly by blocking functional host components from forming working complexes through a dominant-negative interaction^72^.

#### Q73GG5 and Q73HX8

The wMel Q73GG5 and Q73HX8 proteins have inferred ATP-binding and ABC transporter homology and exhibit strong partial alignments with a ABC transporter domain in the *Drosophila eato* gene protein product. The structural alignments have similar TM-scores at 0.86 for Q73GG5 and 0.87 for Q73HX8 (Figure 5A). Q73HX8 has a lower fraction of identical residues at 0.22 compared to the 0.31 of Q73GG5. While host target protein alignment coverage was partial (0.12), a large extent of the wMel query proteins were covered (0.94 and 0.95 for Q73GG5 and Q73HX8, respectively). We found Q73HX8 aligned significantly with 25% of the free living proteomes while Q73GG5 aligned with 90%. This host protein promotes dead cell clearance from the anterior of the oocyte. Knockdowns of the *Eato* gene prevent the clearance resulting in higher nurse cell counts^73^.

**Figure 5.**
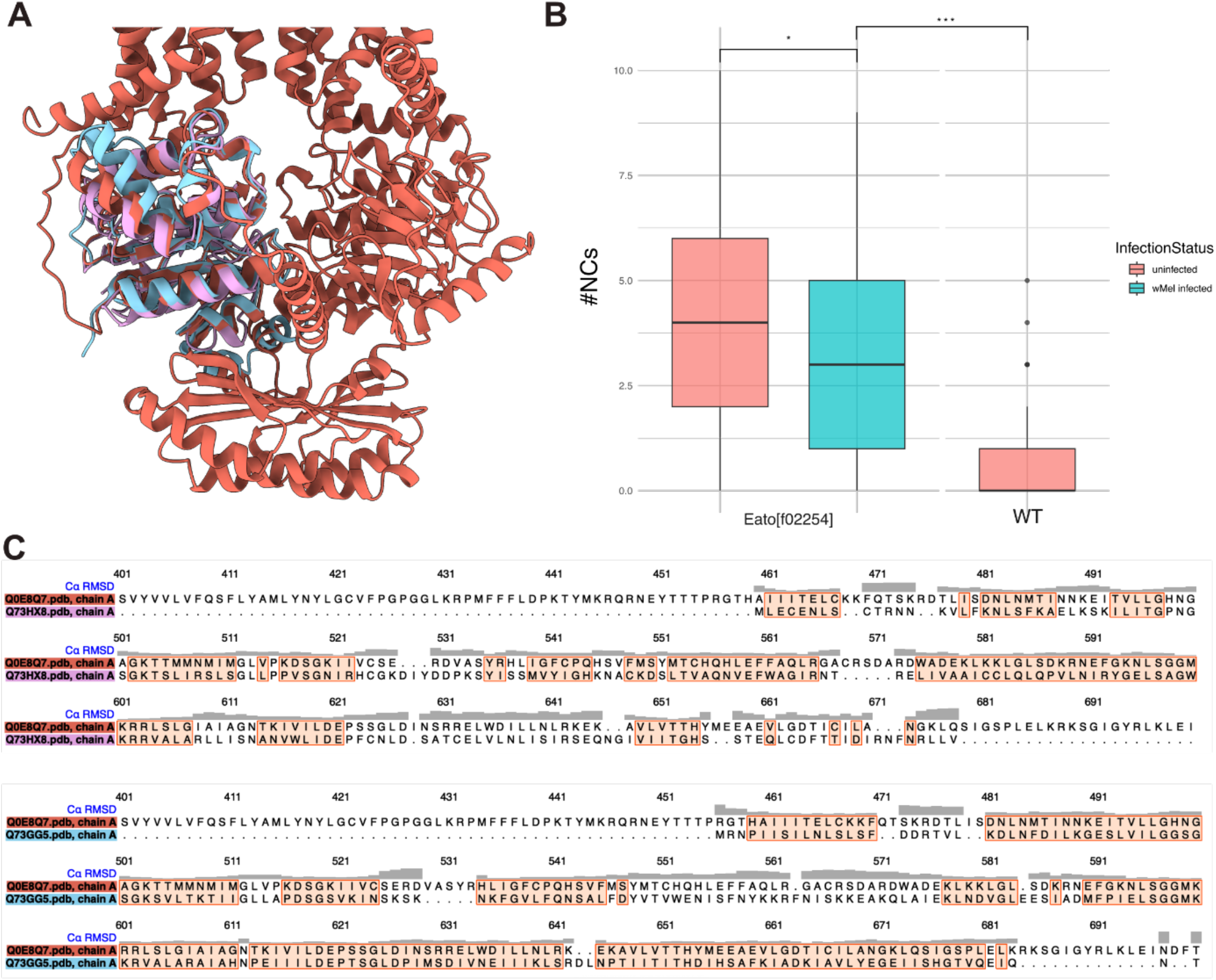
*Eato* Mimic Candidates Overview. **A**. Full structures of Q73GG5 (light blue) and Q73HX8 (pink) aligned with *Drosophila’s Eato* protein. **B**. Nurse cell counts observed in *Eato* knockdown flies both with and without a wMel infection and in the wild type flies expressing *Eato* (Wilcoxon Rank Sum p-value = 0.01167). **C**. Sequence alignment for both candidates with *Eato*.

#### Functional Investigation of *Eato*

Using a nurse cell count assay we investigated the impact of wMel infection on *Eato* function rescue. Our assay demonstrated that *eato* function can be partially rescued by wMel infection (Figure 5B). *Eato* knockouts have 4.17 nurse cells on average (range 0-10) which is substantially more than *Eato*-knockout infected flies (Wilcoxon Rank Sum p-value = 0.01167 average = 3.25, 0-9) and uninfected wildtype (Wilcoxon Rank Sum p-value = 4.91e-16 average = 0.64, 0-4). The full list of cell counts for each sample are provided in Supplemental Table S9. This implies that wMel, which expresses these candidate mimic proteins, is able to partially rescue this phenotype. The structural and functional similarity of the wMel proteins aligned with the *Drosophila Eato* structure suggest that these could be the wMel effectors responsible for the rescue of a component of *D. melanogaster* fertility. However, additional functional evidence is required to demonstrate whether Q73GG5, Q73HX8, or both proteins are specifically responsible for this partial phenotypic rescue effect.

#### P61189

Although filtering protein mimicry candidates by their absence in free-living bacteria was an effective strategy for obtaining unique host-associated candidate genes (Table 1, Figures 1-5), mimics do not need to be unique to host-associated clades to be involved in host-symbiont interactions. Indeed, highly conserved and essential proteins are likely targets for co-evolved interactions due to their stasis over time. Furthermore, targeting these conserved elements may provide a generalistic strategy for associating with different host taxa^74^. To search for examples of highly conserved protein structures found in many other free-living bacteria that may be utilized by intracellular organisms for survival or host manipulation, we investigated alignments that had high TM-scores covering nearly all of the target structure. We looked for lower than expected values for fraction of identical residues and fraction of free-living proteomes aligned for alignments within this region.

The *wMel* Chaperone protein P61189 demonstrates full structural similarity with a host chaperone protein, despite high prevalence among free-living bacteria, suggesting that functional conservation may enable mimicry. P61189 encodes High temperature protein G (HtpG), which has been inferred from sequence homology to be a heat shock protein generally involved in cellular stress response. HtpG aligns strongly with the *Drosophila* cytoplasmic heat shock protein Hsp83, returning a TM-score of 0.9 for an alignment covering 0.941 of the *Drosophila* protein sequence and 0.984 of the wMel sequence (Figure 6). This alignment had a fraction of identical residues of 0.33 and a fraction of free-living proteomes aligned of 0.66. These values were below the averages of the top 100 alignments ranked by TM-score and target coverage which were 0.4 and 0.83 respectively.

**Figure 6.**
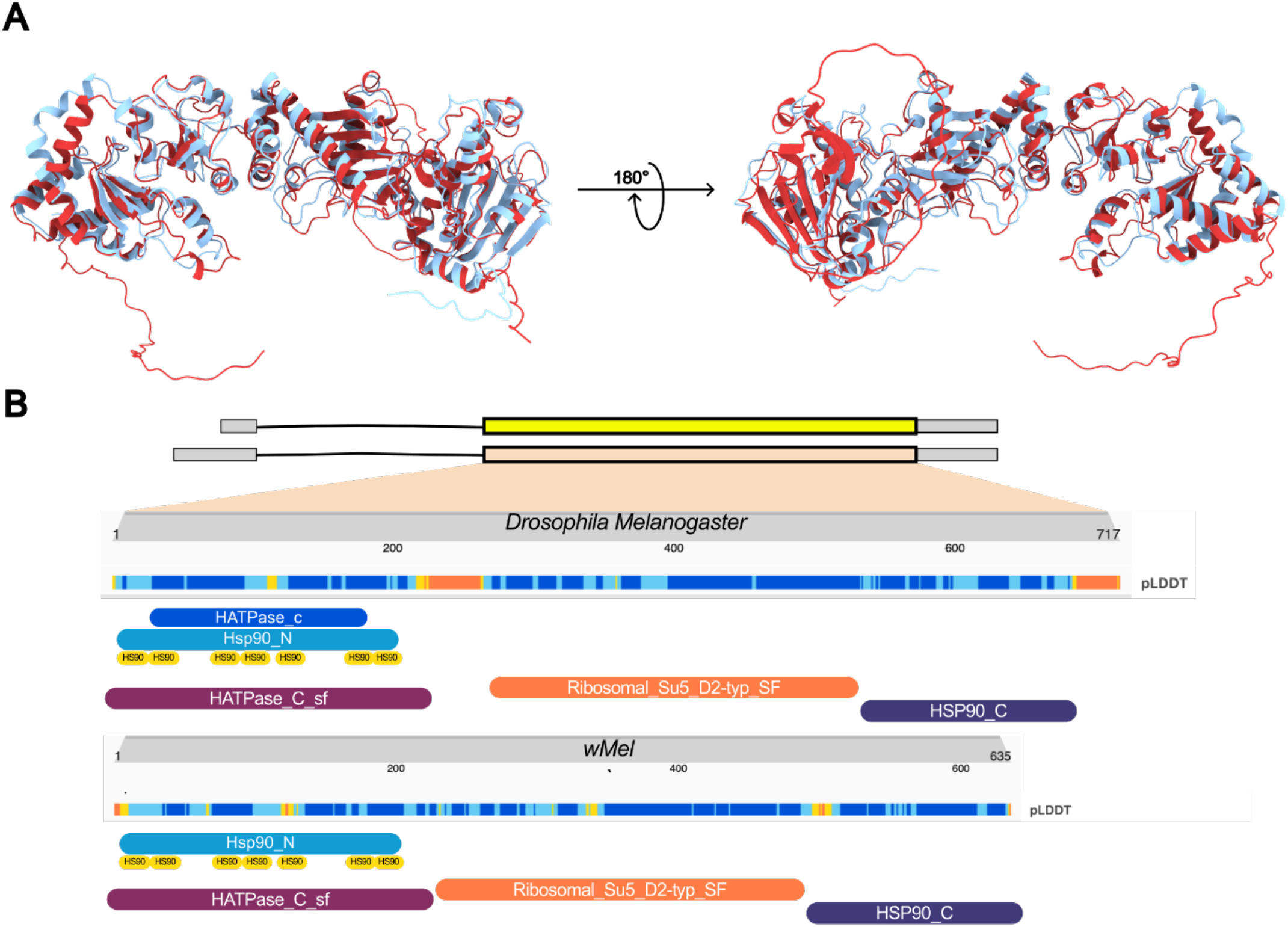
Mimic Candidate P61189. **A**. Structural alignment between the Wolbachia (blue) and Drosophila proteins (red). **B**. Conserved domains across the length of each protein

Phylogenetic inference of HtpG and Hsp83 evolutionary histories reveal that the structural similarity of these proteins is due to deep structural conservation, and not to recent horizontal gene transfer (HGT) (Supplemental Figure S6). As the sequence similarity between wMel and *D. melanogaster* proteins was high compared to other considered mimic candidates (0.33 vs <0.25 for all but one candidate in Table 1), we wanted to test whether the observed structural similarity was due to horizontal gene transfer (HGT) or structural conservation. We collected sequences related to the *Wolbachia* HtpG protein by BLAST^75^ to the non-redundant database^76^, resulting in 425 bacterial, 100 archeal, and 382 eukaryotic taxa. Within the eukaryotic taxa, there were 268 insects, 21 other metazoans, and 93 other non-metazoan eukaryotes. The tree indicates HtpG (P61189) and Hsp83 (P02828) are not close relatives. While both proteins group with their congeners and higher order taxonomic affiliations, e.g., HtpG is an alphaproteobacterial gene and Hsp83 is an insect gene. The midpoint root of the tree suggests that eukaryotic sequences diverged from a shared archaeal or mitochondrial ancestor before the bacterial sequences. Deep within the eukaryotes a duplication-pseudogenization event produced two copies of the protein that are retained in some taxa. Extra copies of the gene appear enriched in symbiont-associated insects, although our dataset is enriched for these groups. Archaeal sequences were scattered throughout the tree and often represented the deepest branching clade in new domain radiations (e.g., at the base of the eukaryotic and pseudomonadota clades). There were several examples of HGT of this protein in other clades (red taxon labels in Figure S6), but the direction and certainty of HGT will need to be confirmed by checking for genome mis-assemblies. Overall, the HtpG/Hsp83 chaperone protein is deeply conserved and archeal taxa appear to have mediated a spread of the chaperone across the tree of life.

Heat shock proteins are evolutionarily conserved between bacteria and eukaryotes, suggesting that these elements may be co-opting ancient housekeeping processes. The D. melanogaster Hsp83 protein is associated with the endoplasmic reticulum (ER) chaperone complex^77^ and wMel is known to alter the abundance and distribution of host ER^78^. The wMel HtpG protein could mediate interactions with the ER and aid in the prevention of host stress responses at a cellular location specific to infection. Furthermore the Hsp83 protein has been shown to act as a specific component of the cap-binding complex where it interacts with translational repressor proteins during oogenesis^79^. The wMel strain is known to play a role in host germ-line development by rescuing the loss of host proteins essential to regulating transcription and translation^80–82^. Specifically, wMel rescues the loss of proteins essential to regulating transcription and translation. Currently the mechanism behind this rescue phenotype is only partially characterized^14^, and HtpG could be an important factor. Hsp83 is also a positive regulator of insulin signaling. It has been found that insulin signaling represses bacterial titer through the TORC1 pathway, potentially through increased autophagy^83^. This goes to suggest a third potential function of *Wolbachia’*s HtpG protein, mimicking Hsp83 to block signaling and repress TORC1 activation.

## Conclusion

The intricate molecular interaction networks of host-microbe systems are an experimentally challenging set of phenomena to research, but computational protein structure prediction and alignment methods can alleviate some of this burden. This approach offers a means of rapidly identifying critical molecular components of host-microbe systems associated with many human diseases and bioengineering applications. Our use of novel structural proteome datasets and tools represents an effective scan of host-microbe systems for structural homology that is suggestive of structural mimicry. Using examples of known structural mimicry we established filters and controls to select structural similarity, applied this structural proteome screen to multiple host-microbe systems, produced controlled structural alignments, and identified candidate proteins that may participate in mimicry. Collectively, our results demonstrate that structure-prediction and alignment approaches are a potentially appealing method for identifying molecular mimicry in host-microbe interactions. Being able to quickly identify potential genes and proteins critical to systems of host control through mimicry can accelerate research looking to perturb or harness host-microbe systems.

We were able to identify a large set of potential mimic candidates in multiple host-microbe systems. The ontology of many of these candidate mimics further corroborated their potential to be actively manipulating host biology in their respective systems. The mimic candidates presented here show strong evidence of mimic-like effectors and suggest mechanisms of host interaction for previously known virulence factors. Our initial screen of the *Drosophila-wMel* system provided strong examples of candidate structural mimics by using stringent filters and selecting proteins with function that can be correlated with *Wolbachia*-host phenotypes. In this system we demonstrated evidence of the predicted mimicry of *Eato*, where we showed *Eato* function in *Drosophila* oogenesis is partially rescued by wMel infection. We also effectively applied the screen to other medically important host microbe systems. In our *Legionella*-human validation system, we identified new potential effectors and candidate mechanisms of interaction for the well-known effector, Mip. Similarly this mimic screen was able to pose new hypotheses of host manipulation by *H. pylori*, which built on the findings of recent biochemical studies looking to identify novel effectors. These results support the use of this framework as a means of both supplementing and suggesting wet-lab experiments for the interrogation of host-microbe systems.

## Supporting information

Supplemental Datafile 4

Supplemental Datafiles 1-3

## Acknowledgements

This work was supported by the NIH (R00GM135583 to SLR; R35GM128932 to RCD; and T32HG012344).

## Code availability

Scripts used for generating datasets and performing analysis are available at: https://github.com/gabepen/mimic_screen

