## Supplementary Information for "A Structural Proteome Screen Identifies Protein Mimicry in Host-Microbe Systems"

### Supplemental Information

#### Tables

#### Table S1. Free Living Database Citations

File included in supplemental data.

#### Table S2. *Legionella* Known Mimic Statistics

| Gene Name | Uniprot ID (LP) | Uniprot ID (Human) | TM-Score | Target Coverage | Query Coverage | Fraction of identical residues | pct_freeliving | average_pLDDT | type |
|---|---|---|---|---|---|---|---|---|---|
| LpdA | Q5ZVD8 | P09622 | 0.957 | 0.927 | 0.991 | 0.413 | 0.981 | 97.173 | Full |
| lpp2176 | Q5ZTI6 | O95470 | 0.710 | 0.944 | 0.991 | 0.320 | 0.411 | 91.494 | Full |
| Lpg1905 | Q5ZUA2 | P49961 | 0.729 | 0.800 | 0.908 | 0.227 | 0.720 | 94.429 | Full |
| LamB/lpg2528 | Q5ZSI9 | Q07837 | 0.493 | 0.621 | 0.848 | 0.111 | 0.794 | 93.003 | Full |
| LamA/lpg1671 | Q5ZUX1 | P04746 | 0.500 | 0.489 | 0.376 | 0.142 | 0.112 | 92.041 | Full |
| RomA | Q5ZWA1 | Q6IQ20 | 0.637 | 0.672 | 0.493 | 0.189 | 0.523 | 68.755 | Partial |
| RalF | Q5ZU58 | Q9NRD0 | 0.759 | 0.564 | 0.525 | 0.274 | 0.000 | 96.381 | Partial |
| LegG1 | Q5ZU32 | O95199 | 0.781 | 0.327 | 0.871 | 0.237 | 0.047 | 92.395 | Partial |
| LegAS4 | Q5ZUS4 | Q9H5I1 | 0.709 | 0.307 | 0.272 | 0.192 | 0.150 | 96.204 | Partial |
| LubX | Q5ZRQ0 | O95155 | 0.710 | 0.271 | 0.450 | 0.268 | 0.000 | 94.837 | Partial |
| Lpg2370 | Q5ZSZ6 | P42356 | 0.400 | 0.116 | 0.836 | 0.115 | 0.047 | 90.977 | Partial |
| LegK1 | Q5ZVF7 | Q9C098 | 0.641 | 0.395 | 0.577 | 0.190 | 0.168 | 88.200 | Partial |

| | | | | | | | | | |
|---|---|---|---|---|---|---|---|---|---|
| LegK4 | Q5ZYU9 | Q13569 | 0.599 | 0.507 | 0.933 | 0.164 | 0.850 | 91.719 | Partial |
| LegK2/Lpg2137 | Q5ZTM4 | O75716 | 0.520 | 0.951 | 0.524 | 0.147 | 0.047 | 79.716 | Partial |
| Lpg2603 | Q5ZSB6 | Q9UQ07 | 0.509 | 0.496 | 0.435 | 0.158 | 0.028 | 90.225 | Partial |

**Table S3. Full Expansive Control Results for *Legionella*-Human.**
File included in supplemental data.

**Table S4. GO Analysis Results for *Legionella*-Human Targets**

File included in supplemental data.

**Table S5. Full Expansive Control Results for *Helicobacter pylori*-Human**

File included in supplemental data.

**Table S6. GO Analysis Results for *Helicobacter pylori*-Human**

File included in supplemental data.

**Table S7. Full Expansive Control Results for *Wolbachia* wMel-*Drosophila.***

File included in supplemental data.

**Table S8. GO Analysis Results for wMel-*Drosophila* Targets**

File included in supplemental data.

**Table S9. Nurse Cell Assay Counts**

File included in supplemental data.

**Data Files**

**Data S1. wMel_GO_analysis_input_IDs.json**

**Data S2. Legionella_GO_analysis_input_IDs.json**

**Data S3. Hpylori_GO_analysis_input_IDs.json**

**Data S4. HtpG-seqdump-all.fasta**

### Figures

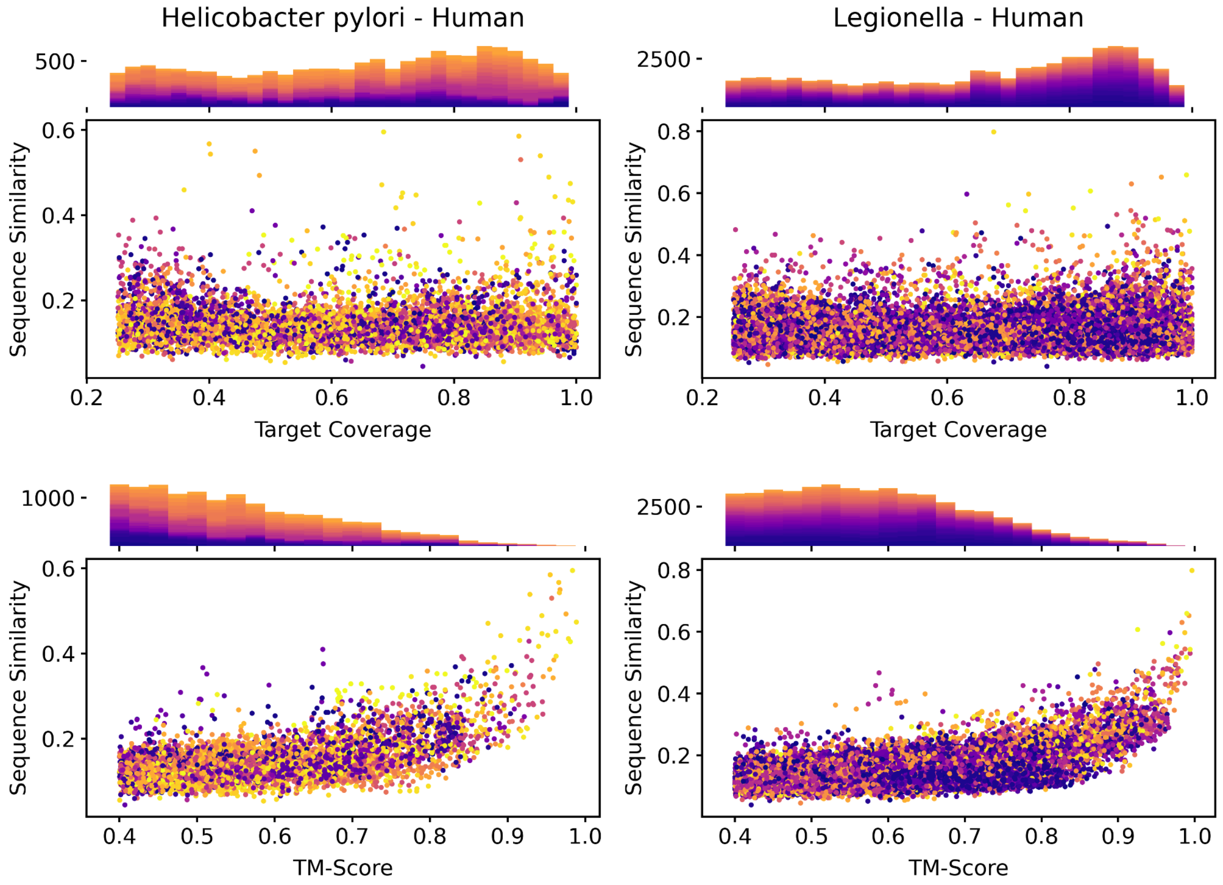


**Figure S1. Expansive Control Alignment Space *Helicobacter pylori*-Human**
*Helicobacter pylori* alignment space compared to the *Legionella* alignment space as described by the expansive control method. Points are colored by fraction of free-living proteomes aligned from the expansive control dataset.

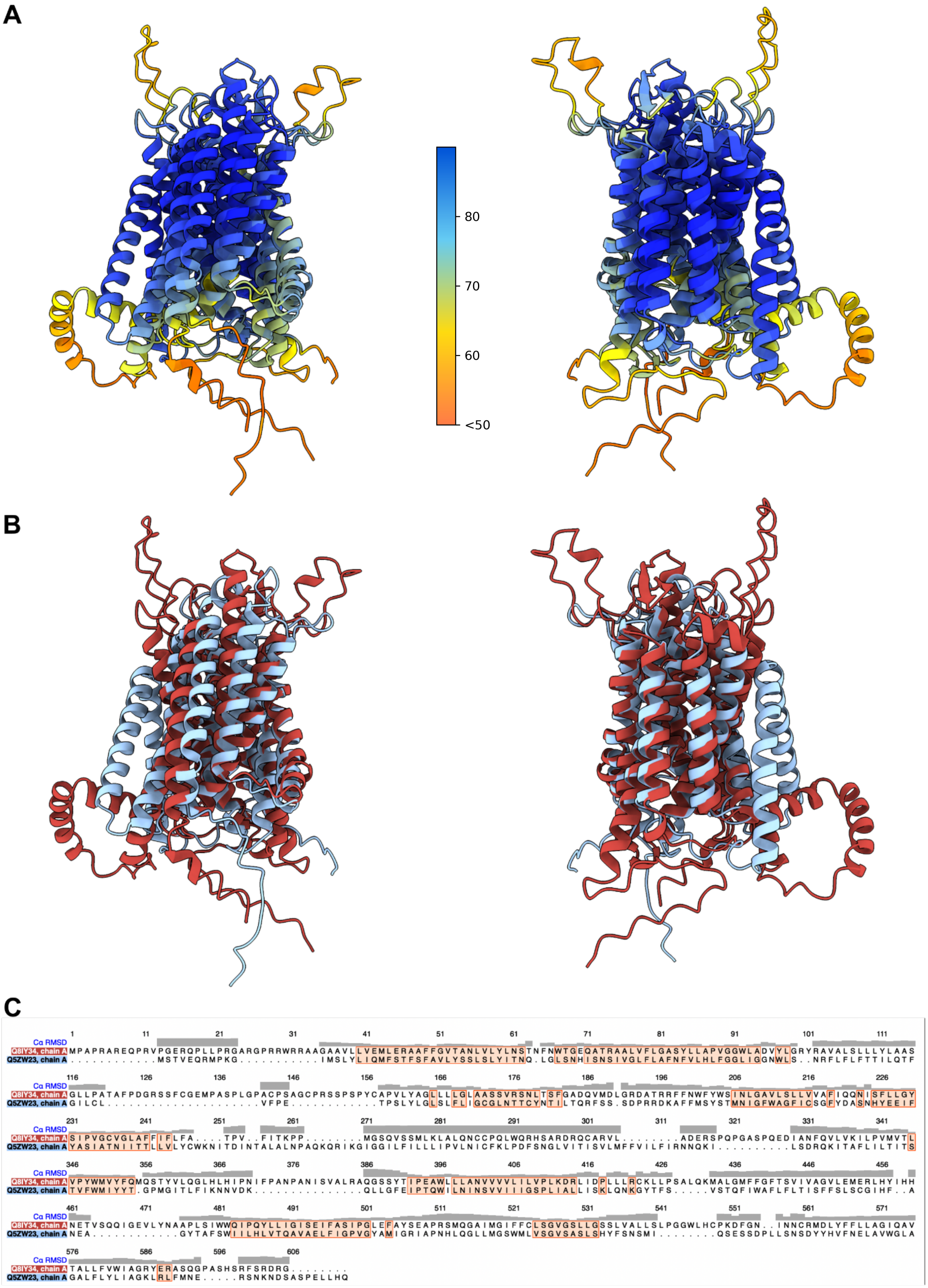

A
80
70
60
<50
B
C
1 11 21 31 41 51 61 71 81 91 101 111
Cα RMSD
Q8IY34, chain A MPAPRAREQPRVPGERQPLLPRGARGPRRWRRAAGAAVLLVEMLERAAFFGVTANLVLYLNSTNFNWTGEQATRAALVFLGASYLLAPVGGWLADVYLGRYRAVALSLLLYLAAS
Q5ZW23, chain A .............MSTVEQRMPKG...........IMSLYLIQMFSTFSFAVLYSSLSLYITNQ.LGLSNHISNSIVGLFLAFNFVLHLFGGLIGGNWLS..NRFLFLFTTILQTF
116 126 136 146 156 166 176 186 196 206 216 226
Cα RMSD
Q8IY34, chain A GLLPATAFPDGRSSFCGEMPASPLGPACPSAGCPRSSPSPYCAPVLYAGLLLLGLAASSVRSNLTSFGADQVMDLGRDATRRFFNWFYWSINLGAVLSLLVVAFIQQNISFLLGY
Q5ZW23, chain A GILCL....................VFPE............TPSLYLGLSLFLIGCGLNTTCYNTILTQRFSS.SDPRRDKAFFMSYSTMNIGFWAGFICSGFYDASNHYEEIF
231 241 251 261 271 281 291 301 311 321 331 341
Cα RMSD
Q8IY34, chain A SIPVGCVGLAFFIFLFA....TPV..FITKPP.........MGSQVSSMLKLALQNCCPQLWQRHSARDRQCARVL.........ADERSPQPGASPQEDIANFQVLVKILPVMVTL
Q5ZW23, chain A YASIATNIITTLLVLYCWKNITDINTALALNPAQKQRIKGIGGILFILLLIPVLNICFKLPDFSNGLVITISVLMFFVILFIRNNQKI...........LSDRQKITAFLILTITS
346 356 366 376 386 396 406 416 426 436 446 456
Cα RMSD
Q8IY34, chain A VPYWMVYFQMQSTYVLQGLHLHIPNIFPANPANISVALRAQGSSYTIPEAWLLLANVVVVLILVPLKDRLIDPLLLRCKLLPSALQKMALGMFFGFTSVIVAGVLEMERLHYIHH
Q5ZW23, chain A TVFWMIYYT.GPMGITLFIKNNVDK...............QLLGFEIPTQWILNINSVVIIIGSPLIALLISKLQNKGYTFS.....VSTQFIWAFLFLTISFFSLSCGIHF..A
461 471 481 491 501 511 521 531 541 551 561 571
Cα RMSD
Q8IY34, chain A NETVSQQIGEVLYNAAPLSIWWQIPQYLLIGISEIFASIPGLEFAYSEAPRSMQGAIMGIFFCLSGVGSLLGSSLVALLSLPGGWLHCPKDFGN..INNCRMDLYFFLLAGIQAV
Q5ZW23, chain A NEA............GYTAFSWIILHLVTQAVAELFIGPVGYAMIGRIAPNHLQGLLMGSWMLVSGVSASLSHYFSNSMI.........QSESSDPLLSNSDYYHVFNELAVWGLA
576 586 596 606
Cα RMSD
Q8IY34, chain A TALLFVWIAGRYERASQGPASHSRFSRDRG........
Q5ZW23, chain A GALFLYLIAGKLRLFMNE.....RSNKNDSASPELLHQ

**Figure S2. Full Q5ZW23 Structure Alignment** Structural alignments between Q5ZW23 (*Legionella pneumophila*) and Q8IY34 (Human). **A.** Structural alignment colored by pLDDT. **B.** Alignment colored by structure. Q5ZW23: light blue, Q8IY34: red. **C.** Sequence alignment and structural RMSD score.

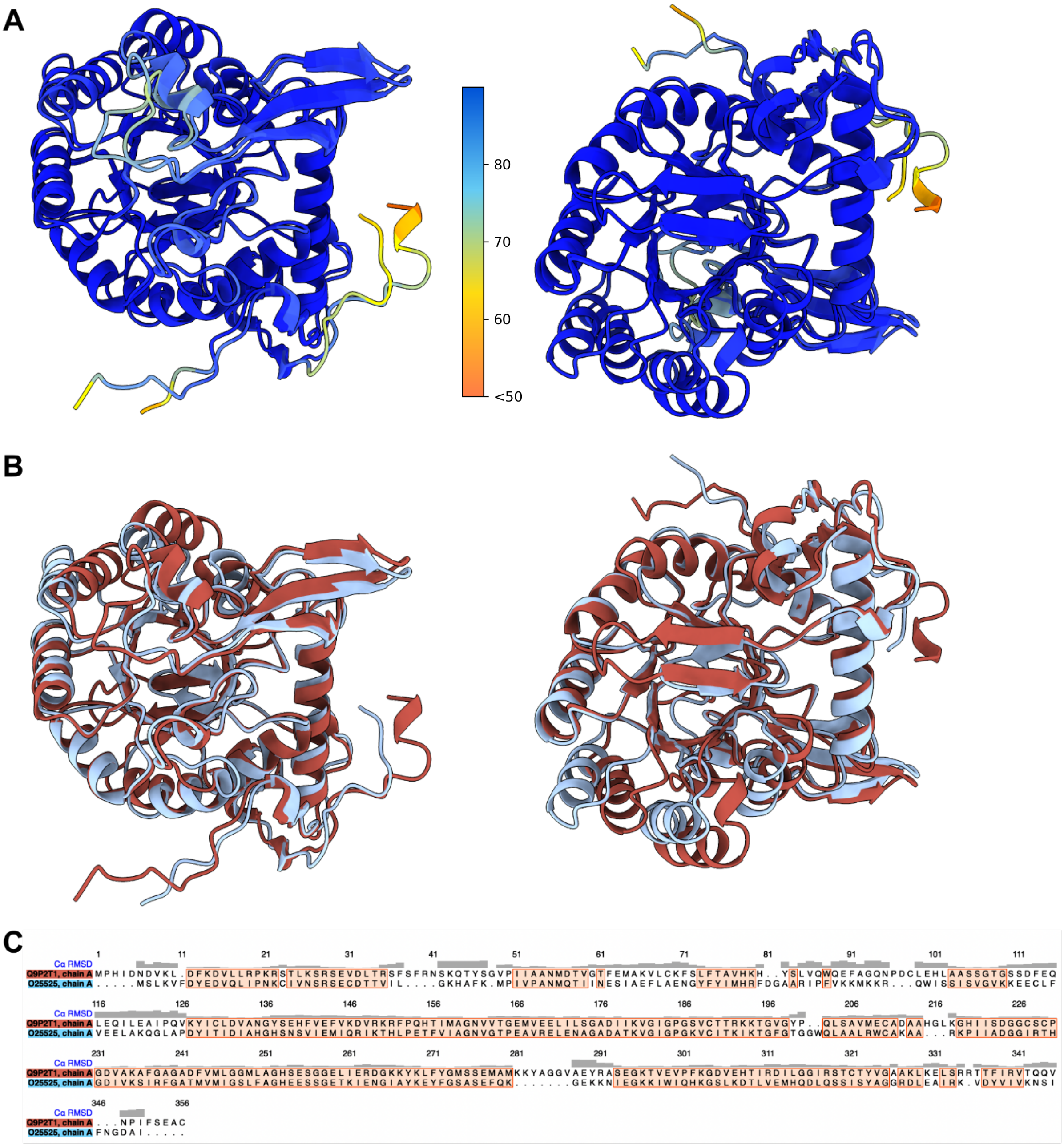


**Figure S3. Full O25525 Structure Alignment** Structural alignments between O25525 (*Helicobacter pylori*) and Q9P2T1 (Human). **A.** Structural alignment colored by pLDDT. **B.** Alignment colored by structure. O25525: light blue, Q9P2T1: red. **C.** Sequence alignment and structural alignment RMSD score.

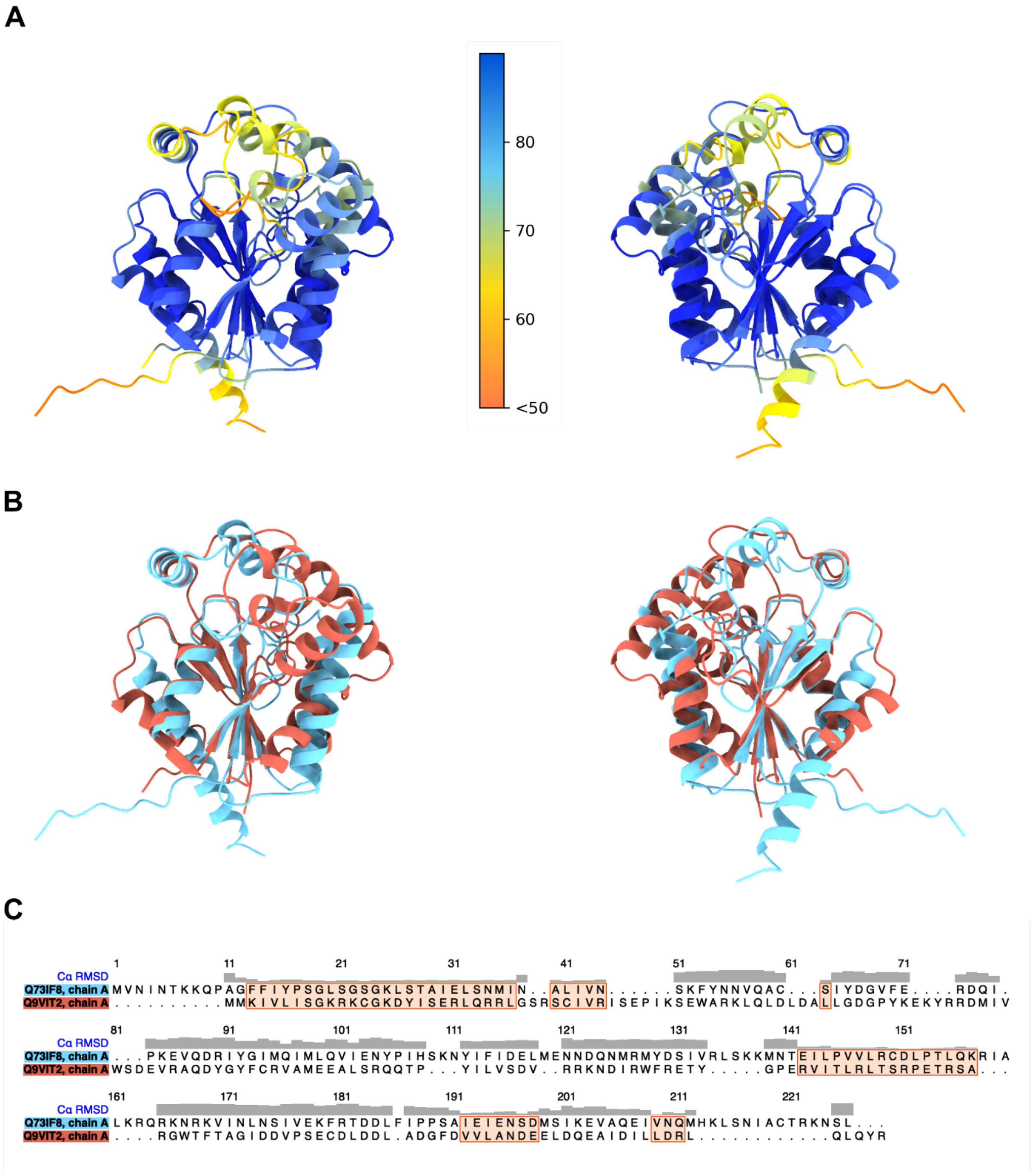


**Figure S4. Full Q73IF8 Structure Alignment** Structural alignments between Q73IF8 (*Wolbachia wMel*) and Q9VIT2 (*Drosophila melanogaster*). **A.** Structural alignment colored by pLDDT. **B.** Alignment colored by structure. Q73IF8: light blue, Q9VIT2: red. **C.** Sequence alignment and structural alignment RMSD score.

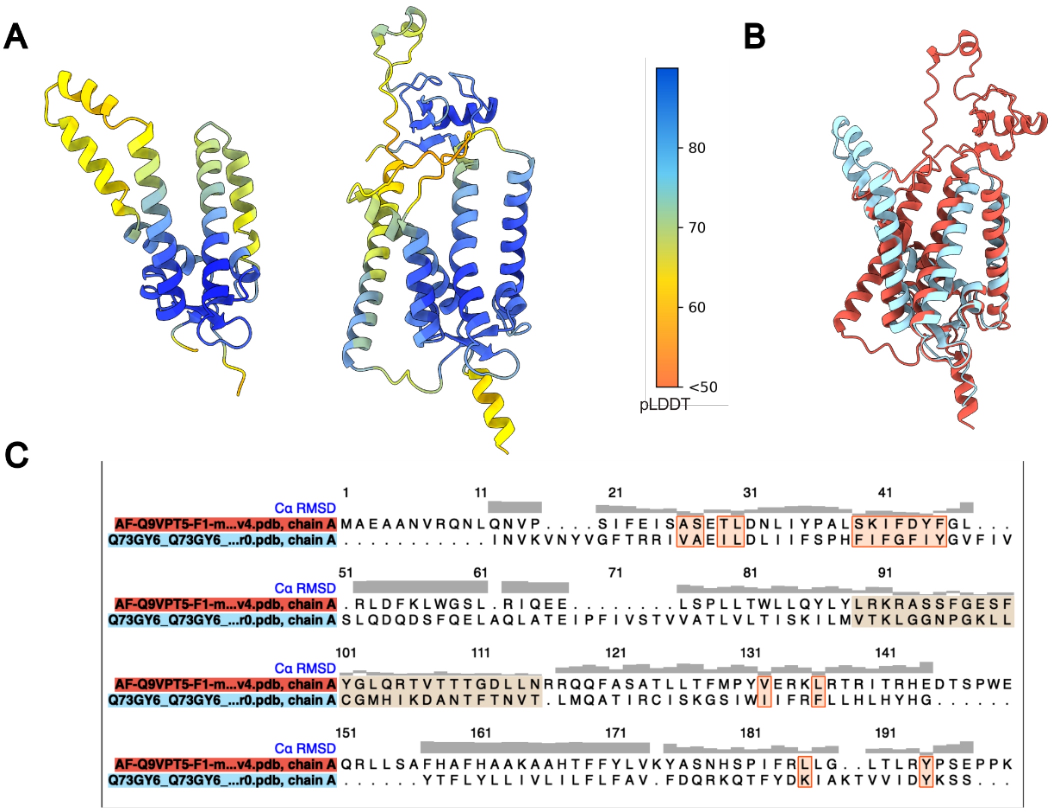


**Figure S5. Mimic Candidate Q73GY6 Overview. A.** Full structures of *w*Mel *Wolbachia* query Q73GY6 (left) and *D. melanogaster* target Q9VPT5, the PEX12 protein (right) colored by pLDDT, showing strong model confidence in the RDD domain region (arrows). **B.** Structural alignment between the wMel and host proteins. **C**. Sequence alignment of the wMel protein and *D. melanogaster's* PEX12 protein, showing low sequence similarity. Structural alignment similarity is shown by the RMSD at the top of the plot.

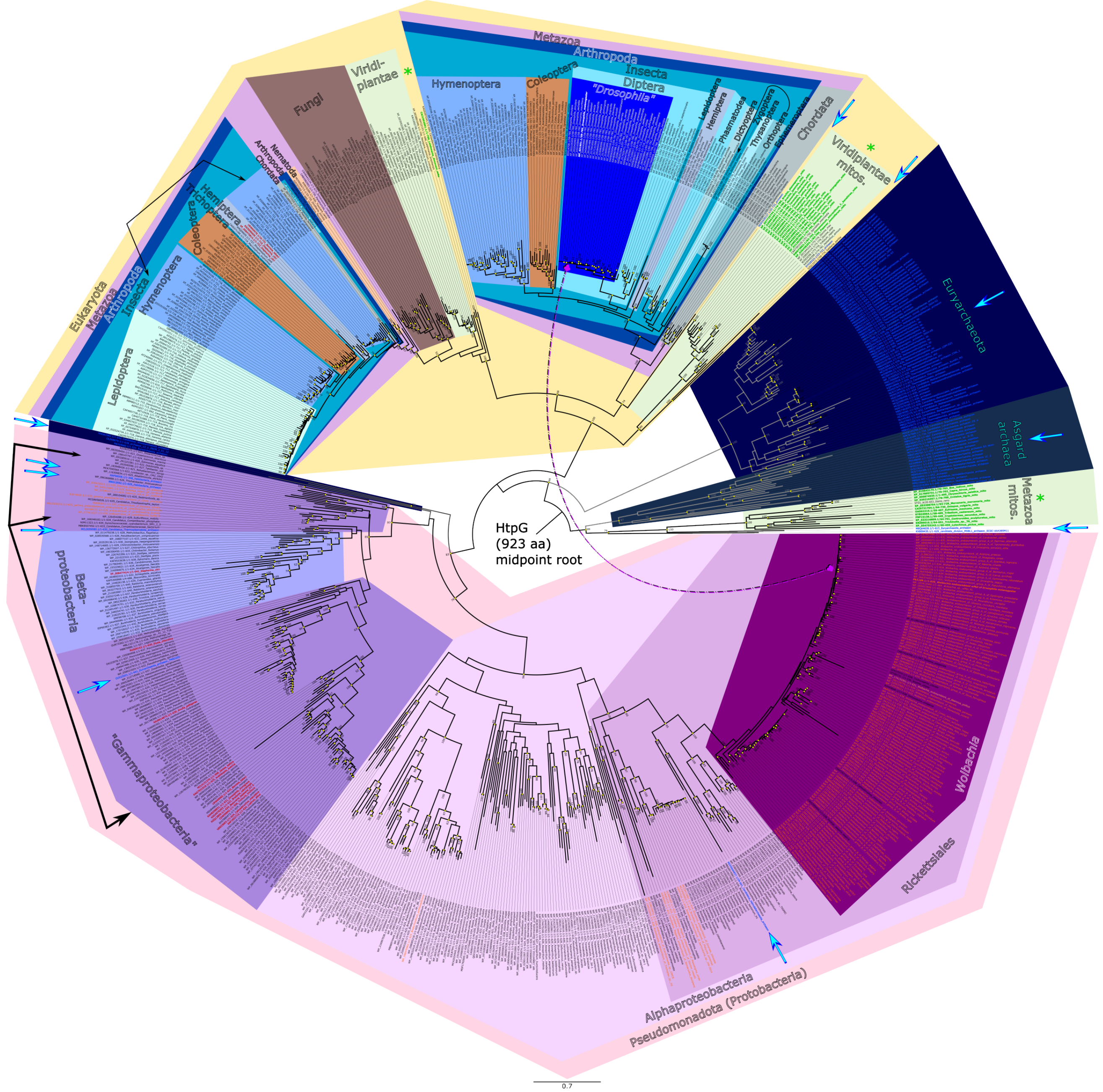


**Figure S6. Mimic Candidate P61189 Phylogenetic Tree.** The *Wolbachia* HtpG and *Drosophila* Hsp83 proteins exhibit highly similar structural similarity, despite deep phylogenetic divergence. Midpoint rooted maximum likelihood tree of 845 chaperone sequences with detectable sequence similarity to *Wolbachia's* HtpG protein, across 923 homologous positions. Numbers at nodes indicate bootstrap support (% of trees supporting consensus topology out of 100 replicates). Values below 50% are omitted. Nodes with >=50% support are denoted with yellow circles. Annotations were added to highlight taxonomic affiliations across the tree and link *Wolbachia* and *Drosophila* proteins (dashed magenta/black line). Dark arrows on the outer perimeter link split taxonomic groupings within a clade. Archaeal taxa are spread throughout the tree, as indicated by the blue (bright/dark) arrows and blue text. Symbiotic taxa are common among the pseudomonadota taxa, and are indicated in orange text. Red taxon names are suspected HGT events (enriched in gammaproteobacteria) or genome misassemblies (e.g., eukaryotic contigs in bacterial assemblies). The black taxon names in the Wolbachia clade are either HGT events (from Wolbachia to the host) or misassemblies (e.g., *Wolbachia* contigs in host assemblies).
